# Ecological drivers of African swine fever virus persistence in wild boar populations: insight for control

**DOI:** 10.1101/2019.12.13.875682

**Authors:** Kim M. Pepin, Andrew J. Golnar, Zaid Abdo, Tomasz Podgórski

**Affiliations:** National Wildlife Research Center, USDA, APHIS, Wildlife Services, Fort Collins, CO; Microbiology, Immunology, and Pathology, Colorado State University, Fort Collins, CO; Mammal Research Institute, Polish Academy of Sciences, Stoczek 1, 17-230 Białowieża, Poland; Department of Game Management and Wildlife Biology, Faculty of Forestry and Wood Sciences, Czech University of Life Sciences, Kamýcká 129, 165 00 Praha 6, Czech Republic

**Keywords:** environmental transmission, carcass, African swine fever, wild boar, persistence, spatial model, transmission, approximate Bayesian computation

## Abstract

Environmental sources of infection can play a primary role in shaping epidemiological dynamics, however the relative impact of environmental transmission on host-pathogen systems is rarely estimated. We developed and fit a spatially-explicit model of African swine fever virus (ASFV) in wild boar to estimate what proportion of carcass-based transmission is contributing to the low-level persistence of ASFV in Eastern European wild boar. Our model was developed based on ecological insight and data from field studies of ASFV and wild boar in Eastern Poland. We predicted that carcass-based transmission would play a substantial role in persistence, especially in low-density host populations where contact rates are low. By fitting the model to outbreak data using Approximate Bayesian Computation, we inferred that between 53 to 66% of transmission events were carcass-based – i.e., transmitted through contact of a live host with a contaminated carcass. Model fitting and sensitivity analyses showed that the frequency of carcass-based transmission increased with decreasing host density, suggesting that management policies should emphasize the removal of carcasses and consider how reductions in host densities may drive carcass-based transmission. Sensitivity analyses also demonstrated that carcass-based transmission is necessary for the autonomous persistence of ASFV under realistic parameters. Autonomous persistence through direct transmission alone required high host densities; otherwise re-introduction of virus periodically was required for persistence when direct transmission probabilities were moderately high. We quantify the relative role of different persistence mechanisms for a low-prevalence disease using readily collected ecological data and viral surveillance data. Understanding how the frequency of different transmission mechanisms vary across host densities can help identify optimal management strategies across changing ecological conditions.

## 1 INTRODUCTION

Understanding mechanisms by which pathogens transmit between hosts is key for defining disease risk and for planning effective control strategies. In addition to direct host-to-host or vector-borne transmission, pathogens can spread through environmental sources, such as through contact with fomites (Allerson et al. 2013), ingestion of contaminated drinking water (Breban et al. 2013, Kraay et al. 2018), contact with contaminated soil (Turner et al. 2014), contact with contaminated carcasses (Chenais et al. 2018), or carcass scavenging (Wille et al. 2016, Brown and Bevins 2018). Environmental sources of infection can promote pathogen persistence by increasing their likelihood of contact with susceptible hosts because many pathogens can remain viable in the environment longer than they can keep a host infectious. For example, epidemiological models demonstrate that pathogens can persist in small populations at very low levels of prevalence when infectious agents remain viable in the environment (Breban et al. 2013). For wildlife populations with seasonally varying densities, environmental sources of infection can ignite seasonal epidemics during low-density periods when susceptible hosts are not frequent enough to continuously maintain pathogen transmission through direct contact (Sauvage et al. 2003). The persistence of infectious agents in environmental reservoirs can enable high pathogen reproductive numbers when epidemic growth rates are low by extending the infectious period beyond the life expectancy of the host (Almberg et al. 2011). In some systems, environmental transmission mechanisms can explain recurrent epidemics (Towers et al. 2018), even at intervals that are longer than demographic cycling (Breban et al. 2013), as well as amplifying rates of inter-population transmission by increasing infectious contact opportunity between groups (Kraay et al. 2018). Theoretical metapopulation modeling has shown that accounting for mechanisms of environmental transmission in addition to routes of direct transmission can lead to qualitatively different disease dynamics and predict different animal movement thresholds for metapopulation decline (Park 2012). Although optimal disease management strategies require knowledge of transmission mechanisms to identify appropriate control points, quantifying the relative role of environmental transmission relative to other transmission mechanisms has been elusive for most host-pathogen systems (but see Towers et al. 2018).

African swine fever (ASF) is a highly virulent disease of swine with devastating consequences for domestic swine industries and food security globally. The virus is known to spread through host-to-host contact, contact with infected carcasses, meat products, fomites, aerosols, the environment, or through tick vectors (Wieland et al. 2011, Costard et al. 2013). Although ASFV persists at low levels among sylvatic hosts in endemic regions of Africa, reports documenting low-level persistence of ASFV in Eastern European wild boar populations in the absence of viral spillover from domestic swine populations or other sources of infection remains inexplicable (Olševskis et al. 2016; Śmietanka et al. 2016; Frant et al. 2017). Considering wild boar contact dead conspecies frequently, infectious carcasses are hypothesized to enable ASFV persistence in wild boar populations (Costard et al. 2013, Probst et al. 2017, Lange and Thulke 2017, EFSA, 2017). Even though the role of carcass-based transmission in ASFV maintenance remains unknown, ASF management strategies in Eastern Europe continue to promote the rapid removal of wild boar carcasses (Costard et al. 2009; Costard et al. 2013, EFSA, 2017).

Carcass-based transmission is a special case of environmental transmission but where the contamination is from biological material. Carcass-based transmission is hypothesized as a potential mechanism allowing low-level persistence because carcasses can remain infectious for long periods of time relative to live infectious hosts. A second hypothesis to explain persistence of ASFV in wild boar populations is that continual introduction from neighboring countries plays a role in persistence. To evaluate these hypotheses, we developed and fit a spatially-explicit mechanistic epidemiological model to spatio-temporal disease surveillance data (Fig. 1) using approximate Bayesian computation (ABC). We estimated the levels of direct transmission, carcass-based transmission, and continued introduction that best explained spatial spreading patterns of ASFV. As a separate objective, we used sensitivity analysis on the transmission mechanism parameters to understand the relationship between host density and the importance of carcass-based transmission. We hypothesized that because wild boar tend not to move very far, carcass-based transmission likely accounts for a substantial amount of overall transmission. Along the same lines, we predicted that the potential role of carcass-based transmission would increase with decreasing host density because at low host densities direct contact would be more limited due to the short-range movement tendencies of wild boar and short infectious period of the virus.

**Fig. 1.**
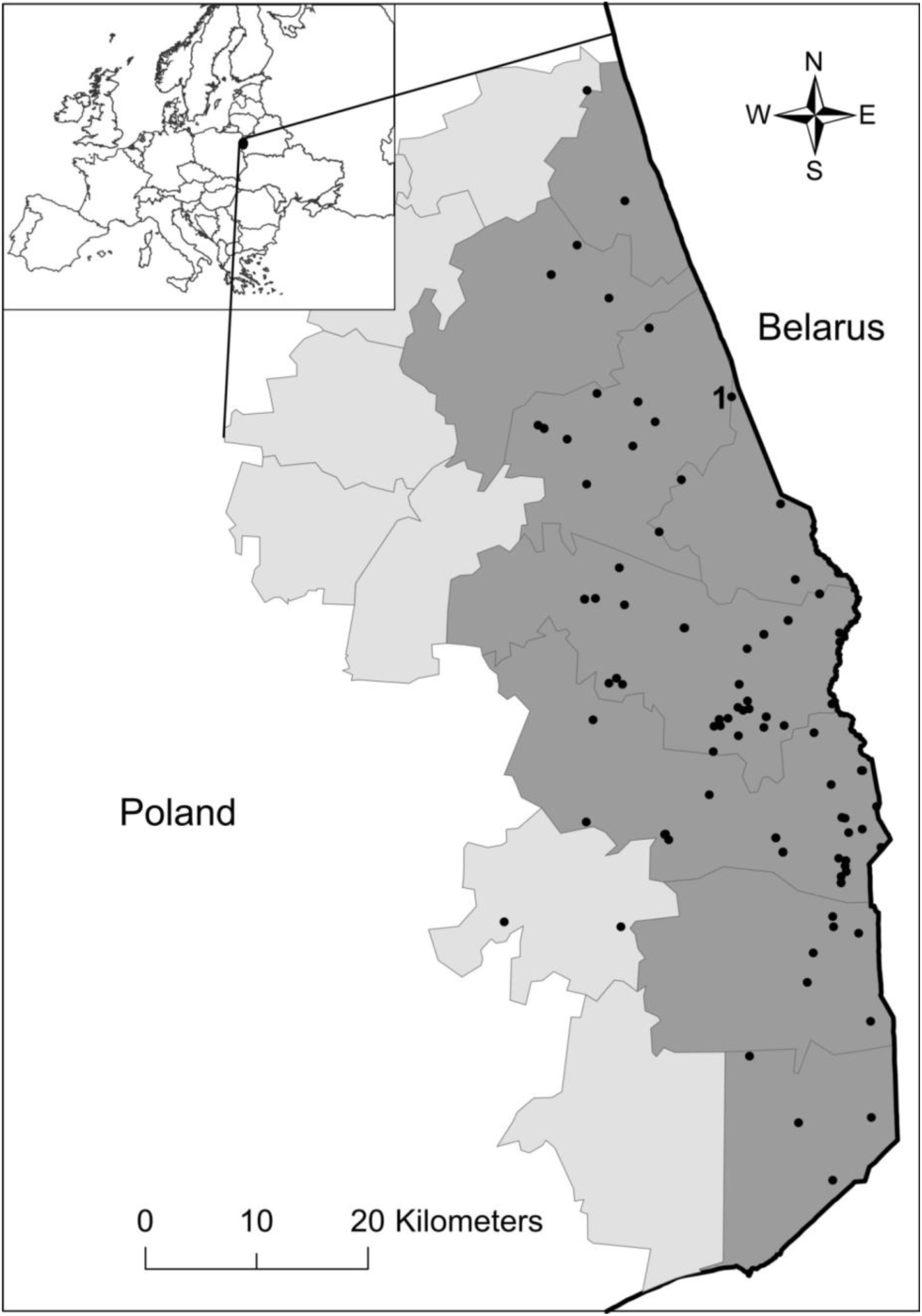
Location of the ASF outbreak in the wild boar population in Poland (2014-2015). Black dots indicate ASF cases in wild boar with the first case labeled. Shaded areas represent administrative districts from which surveillance data used for parameter estimation originated (dark grey - “infected zone” in the text, light grey - “buffer zone”).

## 2 METHODS

### 2.1 African swine fever in Poland

The first wild boar case of ASF in Poland was detected in February 2014 in the north-eastern part of the country (53°19’33”N, 23°45’31”E), less than 1km from the border with Belarus (Fig. 1). Following the first occurrence of ASF in Poland, an intensive surveillance program was implemented in the affected area. The strategy was based on laboratory tests of all wild boar found dead and killed in road accidents (passive surveillance) and all hunted wild boar (active surveillance). A total of 4625 boar were hunted and 271 dead carcasses were sampled in Poland during 2014-2015 (Fig. 1). Samples, collected by veterinary services and hunters, were submitted to the National Reference Laboratory for ASFV diagnostics at the National Veterinary Research Institute in Puławy, Poland. We used surveillance data from 8 administrative districts in Poland where ASFV was detected between 2014 and 2015 (Fig. 1). During this time frame, 139 of 2761 total wild boar samples tested positive for ASFV in a region spanning ∼ 100km along the border (Fig. 1). The furthest case from the border was the 139^th^ case which occurred in late 2015, 34.5 km from the border. Since sampling was strictly linked to hunting activities and the random discovery of carcasses, surveillance data were irregular and seasonally variable. A detailed description of laboratory procedures and tests can be found in Woźniakowski et al. 2016.

### 2.2 Modeling approach

To account for wild boar population dynamics and their impact on ASFV ecology, we developed a spatially-explicit, individual-based model that accounts for variation among individuals in spatial behavior and social interactions (Gabor et al. 1999; Kaminski et al. 2005; Podgórski et al. 2014a, b). We estimated unknown parameters by fitting our model to ASFV surveillance data from Poland using methods of ABC (described below). We estimated rates of new viral introduction and probabilities of direct and carcass-based transmission that best explained the data. Rates of viral introduction from Belarus ranged from a single event to 60 introductions per year (i.e., continuous spillover at the border). Likewise for both direct and carcass-based transmission mechanisms, we considered prior distributions that ranged from 0% transmission probability to 100% transmission probability to neighbors daily. Without prior knowledge on transmission dynamics in this specific system, this design allowed us to estimate the relative contribution of these three persistence mechanisms in explaining the observed surveillance data. Due to extreme computational requirements, sensitivity analyses were completed on a subset of data to broadly evaluate how mechanisms of viral maintenance vary over a range of host population densities. The general schematic for our modeling approach is outlined in Figure S1. All analyses were implemented in Matlab (Version R2016b, The MathWorks, Inc., Natick, Massachusetts, United States). Attributes and model parameters for the model are described below.

### 2.3 Individual-based model

#### Landscape

The landscape was comprised of 5 x 5 km (25 km^2^) grid cells arranged similarly to the outbreak area (Fig. 1, Fig. S2). Grid cells each had a carrying capacity of 0.5-2 boars/km^2^, which controlled heterogeneity in population density across the landscape through density-dependent reproduction. The total landscape size was 120 x 50 km (6000 km^2^).

#### Attributes

Individual-level attributes were monitored and updated at a daily time step. Attributes included disease status, age, sex, unique group identification, dispersal age and distance, status of life, reproduction, age at natural death, *x* coordinate, *y* coordinate, and grid-cell ID. The following disease states were included in the model to track ASFV transmission: susceptible, exposed, infectious (alive), and infectious (dead carcass) (Fig. 2). Sex, dispersal distance, and age at natural death were fixed throughout life but the other attributes changed based on time, age, group size, and grid-cell density. Space was continuous at the individual-level (individual home range centroids were continuous variables assigned to individuals that were located in discrete grid cells). Life status was monitored as alive (i.e., contributing to host population dynamics) or dead (i.e., a carcass on the landscape that can transmit disease but does not move or reproduce). Reproductive status described age-based conception ability, gestation status, and time since last birth for females. Individual attributes were updated based on the following processes: natural mortality, disease transmission, dispersal and social dynamics, surveillance sampling (permanent removal of individuals from the landscape), reproduction.

**Fig. 2.**
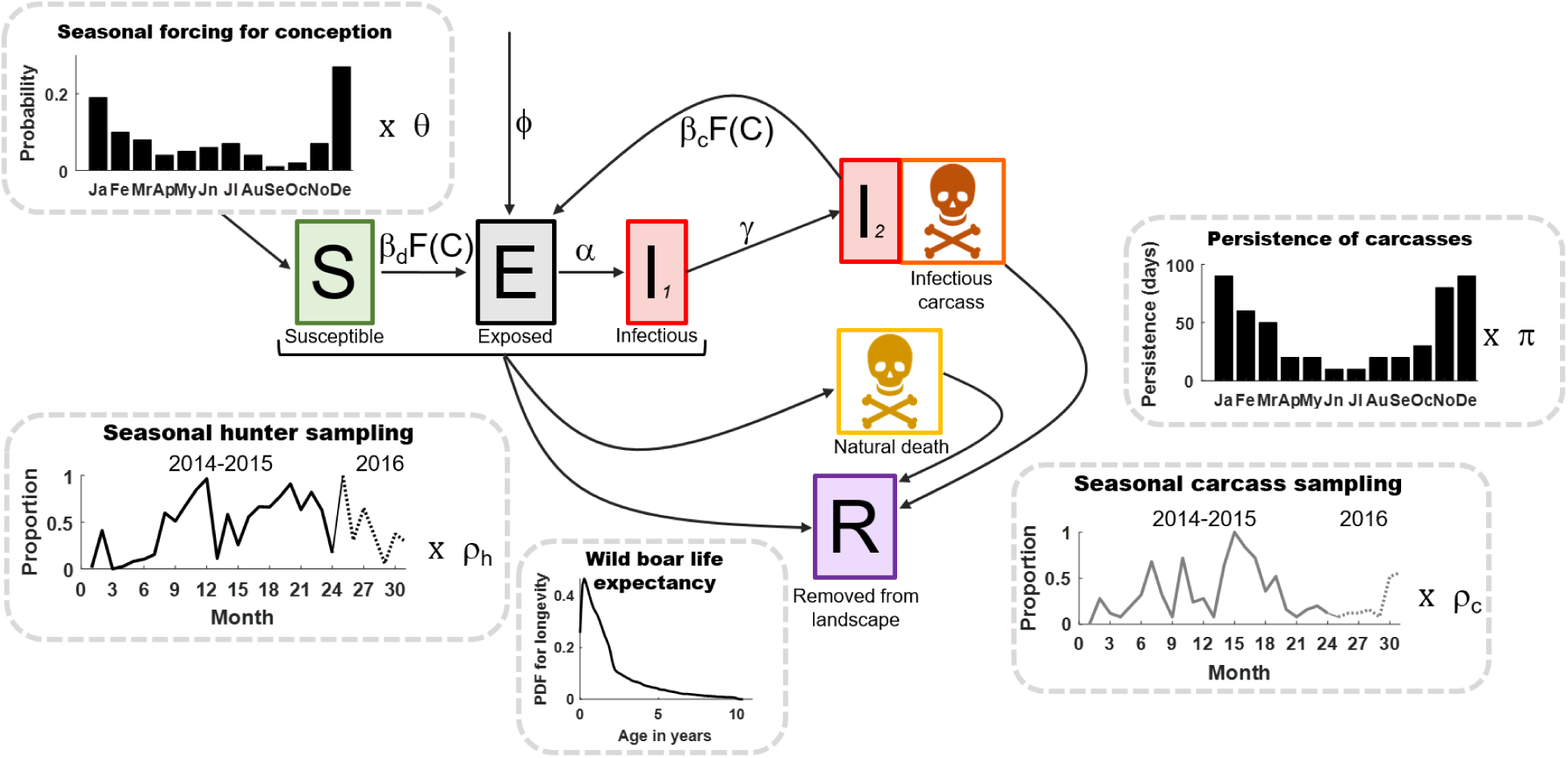
Schematic of disease state transitions and demographic turnover. Seasonal trends in births and carcass persistence are shown in bar plots. Seasonal trends in the intensity of sampling by hunting and carcass sampling are shown in the line plots. There were three mechanisms of mortality: disease-induced (I_2_), natural death, or through hunting. There are two potential routes of transmission: direct (d) or carcass-based (c), which occur via a spatial contact function, F(C), and a transmission probability given contact, β (β_d_ for direct and β_c_ for carcass-based). Persistence of carcasses on the landscape varied seasonally (to reflect weather-based differences in degradation rates) but were the same regardless of the mechanism of death, such that carcasses by all mortality mechanisms had equal probability of being sampled. Seasonal trends in conception probability, carcass persistence, and sampling modes were all multiplied by scaling parameters (θ, π, ρ_h_, ρ_c_) which were estimated. We also allowed for exposed individuals to be introduced along the eastern border at frequency ϕ.

#### Disease dynamics

Disease transmission was modeled using the force of infection equation (FOI; rate at which susceptible individuals become infected) outlined in equation 1, where *x_i,j_* is the distance between the home range centroid of infectious individual *i* (*I_i_*) and susceptible individual *j (S_j_)* (as defined by their x and y coordinates), *a* denotes alive individuals, *b* denotes infectious carcasses, *d* denotes direct transmission, *c* denotes carcass-based transmission, β is the transmission probability that is specific to the transmission mechanism (*d* or *c*). To account for spatial contact behavior in wild boar (Podgórski et al. 2018), transmission probabilities were assumed to decay exponentially with distance according to the rate parameter λ (Table 1). Additionally because wild boar exhibit heterogeneous contact structure due to family grouping (Pepin et al. 2016, Podgórski et al. 2018), probabilities of direct transmission and carcass-based transmission were assumed to be more likely if contact occurred within the same family group (β_wd_ and β_wc_). In equation 1, *w* denotes *I_i_-S_j_* contacts that are within the same family group. Specific parameters are listed in Table 1.

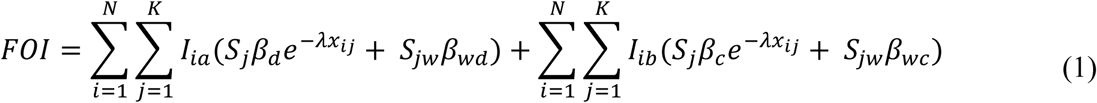

**Table 1.** Model parameters.

| Parameter | Values | Estimated (Y/N) | Source |
| --- | --- | --- | --- |
| **Demographic parameters** |  |  |  |
| Longevity | PDF for longevity in Fig. 2 | N | Jezierski 1977 |
| Daily conception probability per individual | Monthly values in Fig. 2 rescaled to daily (i.e., /30) · $\theta$ | Y ($\theta$) | Ježek et al. 2011; Rosell et al. 2012; Bieber and Ruf 2005; estimated ($\theta$) |
| Litter size | 6 boars | N | Bieber and Ruf 2005; Fruziński 1995; Gethoffer et al. 2007 |
| Age at reproductive maturity (females) | 180 days | N | Gethoffer et al. 2007, Rosell et al. 2012 |
| Minimum time between farrowing and conception | 90 days | N | Barret 1978 |
| Gestation time | 115 days | N | Henry 1968 |
| Age of natal dispersal | ~Poisson(13 months); truncated 10-24 months | N | Podgórski et al. 2014a; Figure S4 |
| Dispersal distance | ~Weibull(2.5,0.5); shown in Figure S3 | N | Keuling et al. 2010; Podgórski et al. 2014a, Prevot and Licoppe 2013; |
|  |  | N |  |
| Maximum group size | 10 | N | Podgórski et al. 2014a |
| **Epidemiological parameters** |  |  |  |
| Incubation period | ~Poisson(4 days), truncated at 1 | N | Blome et al. 2013;Gallardo et al. 2015 |
| Infectious period | ~Poisson(5 days), truncated at 1 | N | Blome et al. 2013, Gallardo et al. 2015 |
| Disease-induced mortality (DIM) | 100% | N | Blome et al. 2013,Gallardo et al. 2015 |
| Contact probability given distance | $e^{-\lambda x_{ij}}$ | Y | estimated |
| Direct transmission probability | $\beta_d$ | Y | estimated |
| Carcass-based transmission probability | $\beta_c$ | Y | estimated |
| Direct transmission probability for contact pairs in the same family group | $\beta_{wd}$ | Y | estimated |
|  |  | Y |  |
| Carcass-based transmission probability for contact pairs in the same family group | $\beta_{wc}$ | Y | estimated |
|  |  | Y |  |
| Persistence of carcasses ($\pi$) | $\pi$ · data in Fig. 2 | Y | estimated ; Selva et al. 2005; N. Selva pers. comm. |
|  |  | Y |  |
| Frequency of spillover ($\phi$) | $\phi$ | Y | estimated |
| **Surveillance parameters** |  |  |  |
| Seasonal trend in sampling | Fig. S4 | N | Unpublished data of the National Veterinary Research Institute, Poland |
| Number of hunter surveillance samples/ day | $\rho_h$ · Fig. S4 | Y | estimated |
| Number of carcass surveillance samples / day | $\rho_c$ · Fig. S4 | | estimated |

Numerous studies demonstrate that animals in the family Suidae are extremely susceptible to multiple strains of ASFV with nearly 100% of domestic pigs and wild boar succumbing to disease within approximately 8-20 days post exposure (Blome et al. 2012, Gallardo et al. 2017). Infection can also present in a chronic form with viral shedding lasting more than 6 weeks, however, attenuated viral strains that promote chronic infections have never been reported in the Eastern European region (Sanchez-Vizcaino et al. 2015). Considering host competence in surviving wild boar is largely unknown, and only a small fraction of individuals are likely to survive viral exposure, we assume ASF is 100% lethal in wild boar. Periods of latent and infectious disease in live hosts were Poisson random variables (Table 1). We tied viral persistence time in carcasses to carcass decay rates to give uninfected and infected carcasses the same opportunity to be sampled (and we could not find data to suggest otherwise). Therefore, infected carcasses were assumed to remain infectious for the entire duration they persisted in the environment. The infectious period of carcasses were assumed to vary seasonally based on field measures of carcass persistence in Eastern Poland (Table 1, Fig. 2).

#### Social structure and dispersal

Social structure of wild boar is based on cohesive, matrilineal groups, composed of a few subadult and adult females and their offspring (Gabor et al. 1999; Kaminski et al. 2005; Podgórski et al. 2014a, b). Studies demonstrate that the frequency of direct contacts is much higher among individuals within than between groups (Podgorski et al. 2014a; Pepin et al. 2016). Further, social groups may temporarily break, reform, or exchange individuals (Gabor et al. 1999; Poteaux et al. 2009) but group members usually form stable and long-lasting relationships (Podgórski et al. 2014a). Thus, social and spatial movement behavior can constrain wild boar contact and modulate the spread of infectious diseases (Loehle 1995). We accounted for the effects of social structure as described in Figure S3. Females and immatures occurred in family groups. Members of the same family group had the same home range centroid. Adult males were independent (not part of a group, each having a unique home range centroid). Social structure was dynamic – family groups that became too large (according to a maximum group size parameter; Table 1), split in half and one group dispersed (Fig. S3). Likewise, because adult females are rarely found alone (Kaminski et al. 2005, Podgórski et al. 2014a), independent females were joined to the nearest family group that was below capacity (Table 1; Maximum group size). Dispersal caused permanent relocation of the home range centroid (*x* and *y* coordinates). Dispersal distances were chosen at random from a Weibull distribution (Table 1; Figure S3). In addition to dispersal due to social structuring, natal dispersal also occurred, but only once at a randomly selected age (Table 1) assigned at birth. Males dispersed independently and females dispersed with their sisters. Although the dispersal of young wild boar leaving their maternal groups is the main source of long-distance movement, the majority of individuals disperse in relatively short distances (1-3 km diameter) from an average home range (<5 km^2^) and longer dispersals (5-30km) are less common (Truve and Lemel 2003; Keuling et al. 2010; Prevot and Licoppe 2013; Podgórski et al. 2014b; Kay et al 2017).

The dispersal process (natal or other relocation) was: 1) for each 45 degree angle from the home range centroid, a new possible set of [x,y] coordinates was obtained using the dispersal distance value assigned at random to the group (Table 1; i.e. x = distance x cos(angle) + current x coordinate, y = distance x sin(angle) + current y coordinate). If at least one of these potential locations were valid (i.e., in a grid cell with fewer boars than the carrying capacity or a location off the grid), then a valid potential location was chosen at random and boar(s) were relocated there. Boars that traveled off the grid were lost permanently. If there were no valid locations, the distance value was doubled and the process repeated until a valid location was obtained.

#### Birth and death parameters

Boar conception occurred randomly in reproductively active females based on a seasonally varying conception probability (Table 1; Fig. 2). Pregnant females gave birth to 6 offspring (3 male, 3 female) after a gestation period of 115 days (Table 1). Following birth there was a fixed lag of 3 months before the possibility of conceiving again (Table 1). Thus, the maximum number of litters per year was 2. Net population growth rate was controlled by multiplying the seasonal trends in conception probability by a scaling parameter (Ɵ). The full range of the prior distribution of Ɵ allowed net population growth rates to range between 1.3 and 2.3 for population densities at 10% of the carrying capacity, consistent with Bieber and Ruf [2005]. Conception probability was density dependent such that conception did not occur in individuals in grid cells that were already at carrying capacity. The population-level host demographic dynamics were similar to a logistic model (Pepin et al. 2017).

Sources of mortality included natural mortality, disease-induced mortality, and hunter harvests (described below). For natural mortality, each individual was assigned a longevity at birth based on wild boar life expectancy (Table 1; Fig. 2).

#### Initial conditions and demographic burn-in

Populations were initialized as follows. A matrix with the number of rows equivalent to the desired population size was created. Each individual (row) was assigned attributes at random (Table 1). For males whose age was beyond dispersal age, dispersal status was recorded as completed. All females and males less than dispersal age were divided into group sizes that were ¼ of the maximum sounder size (plus one smaller group of remaining individuals if applicable). Each individual or group was assigned to a grid cell ID chosen at random (the algorithm ensured that unoccupied grid cells were selected first). Within each grid cell, the individual or group was given [x,y] coordinates selected at random. After the population was initialized, population dynamics were allowed to occur for 10 years. The population at the end of the 10 years was used as the starting point for all simulation conditions with disease transmission.

#### Surveillance parameters

Because hunter harvests made up most of the sampling (94.5%) and hunter harvesting is thought to be a primary regulator of population density (Keuling et al. 2013, Massei et al. 2015), we included it as a source of mortality in our model in addition to using it as our observation model. In 2014, average wild boar densities were estimated at 1.5-2.5 boar/km^2^, locally ranging from 0.5-1 boar/km^2^ to 3-5 boar/km^2^ (Regional Directorate of State Forests, Białystok, Poland). However, because we had no data on how the absolute number of boar sampled related to the underlying density, we added parameters ρ_h_ and ρ_c_ to scale the absolute numbers of boar sampled up or down (Table 1). First we calculated the relative number of boar sampled daily by each surveillance method (number sampled on day t/maximum ever sampled on any given day) to produce seasonal trends in the proportion of the population sampled (Fig. 2; Fig. S4). Next, we multiplied the seasonal trend data for each surveillance method by the scaling factors (ρ_h_ and ρ_c_, Table 1; Fig. S4) to determine the daily proportion of boar that would be sampled by hunter harvesting or dead carcasses. The product of the trend data and the scaling factor can be thought of as a daily detection probability. We assumed that boar < 6 months of age would not be hunted (typically not targeted by hunters) and that boar < 3 months of age would not be sampled by the dead carcass method (because they are unlikely to be found). We recorded the disease status for all boar that were sampled and then immediately removed them from the landscape permanently.

### 2.4 Approximate Bayesian Computation

We estimated the unknown parameters using ABC with rejection sampling. Estimated parameters are indicated in Table 1.

Approximate Bayesian computation selects parameter sets for the posterior distribution using distance metrics (difference between model predictions and the observed data), a measure of how well a model parameter set approximates target patterns in the observed data. We used three distance metrics concurrently; the sum for each of: 1) monthly cases from carcasses, 2) monthly cases from hunter-harvest sampling, and 3) monthly maximum distance from the border. Distance metric tolerance values were 48 for monthly cases from carcasses, 24 for monthly cases from hunter-harvest samples, and 120 for maximum distance from the border. Parameter sets with outcomes lower than these values for all 3 metrics comprised the posterior distribution. This allowed average error rates of 2 (carcass) and 1 (hunter harvest) cases, and 5 km from the border per month on average. We chose tolerance values based on what we believed to be an acceptable level of error for planning control strategies and risk assessment. Also, more stringent error rates would require restrictively large computational resources unless prior distributions are more informed.

We fit the model to 4 different landscapes separately: ‘patchy’ including high (2 boar / km^2^) and low (0.5 boar / km^2^) density patches guided by the location of cases in the real data (average density ∼1 boar/ km^2^); and ‘homogenous’ landscapes with densities of 1, 1.5, and 2 boar / km^2^. This design evaluated whether observed outbreak patterns could have arisen from the underlying distribution of boar density being higher in patches where the disease was observed relative to other patches, as opposed to alternative mechanisms such as the surveillance patterns.

#### Prior distributions

Each parameter had a uniform prior distribution as follows: frequency of introduction ϕ∼Unif(0,60), β_d_∼Unif(0.0001,1),β_c_∼Unif(0.0001,0.99), ρ_h_∼Unif(0.0005,0.1), ρ_c_∼Unif(0.0005,0.8), π∼Unif(0.1,1.5), θ∼Unif(0.5,6), λ ∼ Unif(0.1, 2.5), β_wd_∼Unif(0.01,1), and β_wc_∼Unif(0.001,1). Prior distribution ranges were informed by movement and contact data (Podgórski et al. 2013, Pepin et al. 2016, Kay et al. 2017, Podgórski et al. 2018). As part of the parameter generation process we implemented the following constraints for each parameter set: β_d_>β_c_, β_wd_>β_d_, β_wc_>β_c_; to further inform prior distributions with biologically realistic knowledge. To sample across parameter space efficiently we used a Latin hypercube algorithm to generate 979,592 parameter sets and then ran the model twice on each parameter set (for a total of 1,959,184 iterations; or 2 chains of 979,592). β_d_, β_c_, and ρ_c_ were sampled on a log_e_ scale. Because the epidemiological model was time-intensive we used a two-tiered approach to evaluating parameter sets. First, simulations were terminated early if the trajectory was unrealistic – specific criteria were: 1) the landscape-wide host density dropped below 20% of the initial density, 2) more than 150 new cases occurred per day; 3) there were no new cases sampled by either type of surveillance method in the past 6 months, or 4) the total number of cases sampled by both methods of surveillance totaled more than 300 (more than double the actual number). We then only considered parameter sets for which the simulation reached the end of the two-year time frame. For this reduced set, the posterior distributions consisted of all unique parameter sets (considering both chains) that were within the absolute distance of three metrics: the sum of absolute differences between observed and simulated data for monthly positive samples from live and dead animals (considered separately), and the maximum monthly distance of cases from the border.

#### Goodness-of-fit

To determine the ‘best’ landscape model we ranked models from the different landscapes based on their distance metrics (where minimum values are best) and the R^2^ values (squared correlation coefficient of the observed and predicted data) for the observed versus the predicted monthly cases and monthly distance from the border (Table 1). To calculate the R^2^ values, for each landscape we conducted 1000 simulations using random draws from the posterior distributions of parameters and calculated the R^2^ value for each simulation. We then took the mean of the 1000 R^2^ values for each metric (monthly cases and distance) to represent the overall R^2^ values for the metrics of a particular model. As another measure of predictive ability, we tested the ability of our models to forecast ASF dynamics by using the parameters estimated from fits to the 2014-2015 data to predict the first 7 months of 2016 (Jan.-Jul.). We then compared the R^2^ values for the in-sample predictions relative to the full set of predictions (Table 1).

### 2.5 Sensitivity analyses

Sensitivity analyses were conducted on homogenous landscapes varying in density from 1-4 boar / km^2^, reflecting the observed densities of wild boar in Eastern Europe (Melis et al. 2006). We completed a full factorial sensitivity analysis to assess how ASF persistence and transmission dynamics respond to changes in ϕ, β_d_, and β_c_. Transmission parameters β_d_, and β_c_ varied from no transmission (0.0001) to high levels of transmission (0.3) and ϕ was varied from 1 introduction to 50 introductions per year. All other parameters were fixed with a parameter set from the posterior distribution of the patchy landscape model. Sensitivity analyses were completed using 3 different 50 km x 50 km landscapes that varied in host density (1-4 boars/km^2)^. The index case occurred in grid cell 50 (middle of the most right side column of grid cells) on day 30 (same day of introduction in the ABC analyses). All runs were conducted for 2 years. We ran 100 replicate simulations for each set of conditions. We recorded all cases (true behavior), but included host mortality due to hunting and removal of dead carcasses due to surveillance sampling. We recorded the following output: 1) persistence probability (proportion of 100 simulations where at least one case occurs in the last week of the two year period after only a single introduction at the start), and 2) the proportion of transmission events that were from direct and carcass-based transmission. The latter output was obtained by recording the proportion of transmission events that were direct transmission for each day and taking the median value over time, considering only days where at least one transmission event occurred. We modeled the outputs using generalized linear models using appropriate distributions and/or data transformations for each of the 4 response variables and including the transmission probability parameters and introduction frequency as covariates, and all interactions. The purpose of these models was simply to interpolate the relationships at a higher resolution within the range of values used in the simulations. For modeling persistence probability, we also included up to 4^th^ order interactions because the relationships were highly non-linear and thus these were important for accurately interpolating the relationships.

## 3 RESULTS

### 3.1 Model fit

Despite high uncertainty in several estimated parameters, the models captured the general trends in the surveillance data well (Fig. 3). All models captured monthly cases better than monthly maximum distance from the border (Table 2, Fig. 3). Relative to the observed data, the model predicted higher incidence during months 14-16 (Feb.-Apr. of the second year which included a birth pulse) and lower incidence than the observed data in months 5-7 (May-Jul. of the first year, which included a period of abnormally low surveillance) (Fig. 3a, Fig. S4). The average prevalence observed through surveillance in the model tracked the magnitude of true sample prevalence for both hunter-harvest and carcass surveillance samples (Fig. S5). Models fit on homogenous landscapes of host density did not capture spatial spreading rates as well as the patchy landscapes that included high-density patches (2 boar / km^2^; average 1 boar / km^2^; Table 2).

**Fig. 3.**
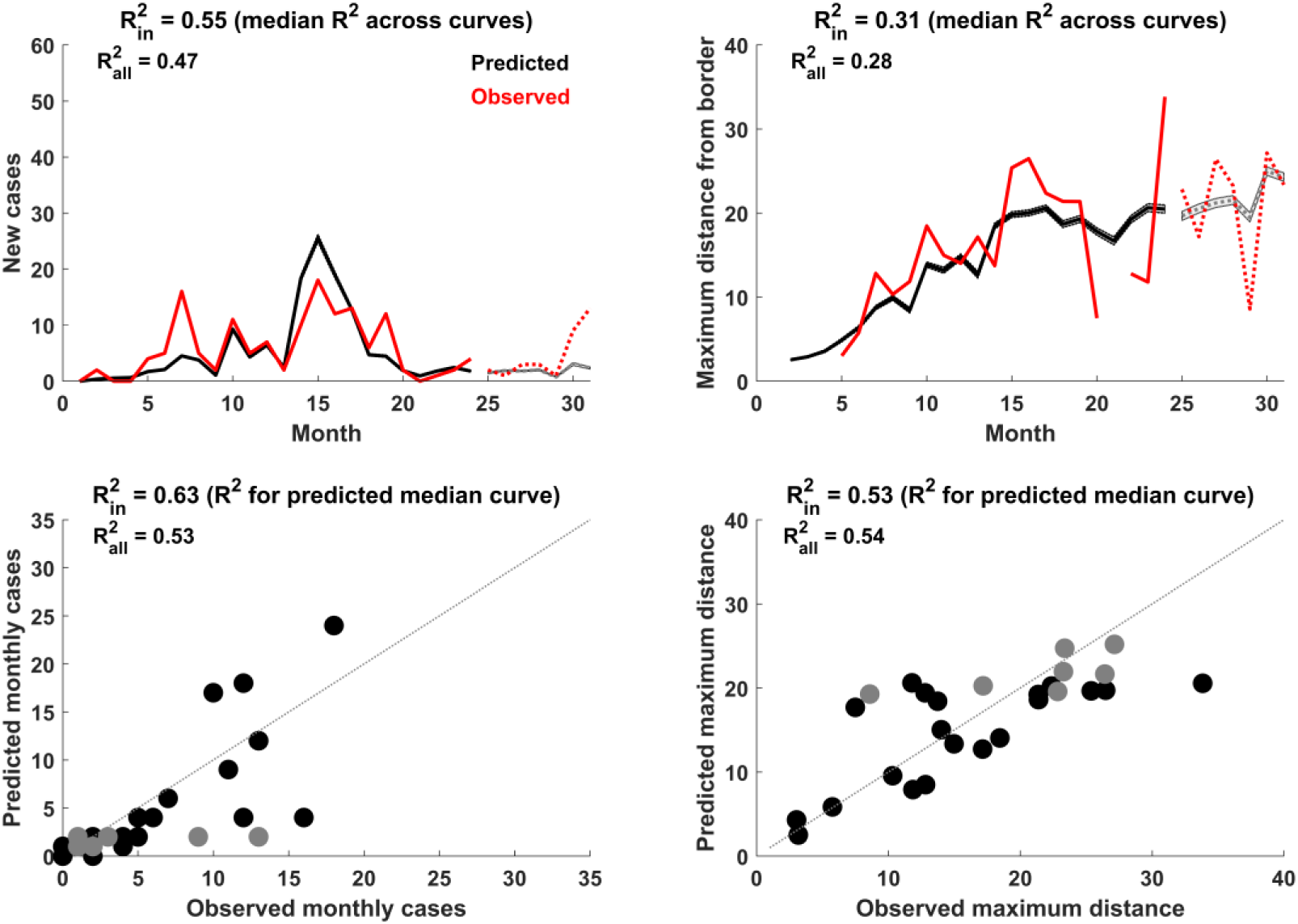
Model fit (patchy landscape). Trajectory of new cases (A) and maximum distance from the border (B) for observed (red) and predicted (black). Shaded areas indicate 95% prediction intervals from 1000 simulations from the posterior distributions of parameters. Solid lines indicate the data that were used for parameter estimation whereas dotted lines show the out-of-sample predictions. C and D show the observed versus predicted points (where each point is the median across all simulations at each time step) for monthly cases (C) and maximum distance from the border (D). In-sample points (2014-2015) are in black, out-of-sample points (2016) are in grey. The grey dotted line indicates the expected fit of the points (1:1 ratio of observed and predicted points).

**Table 2.**
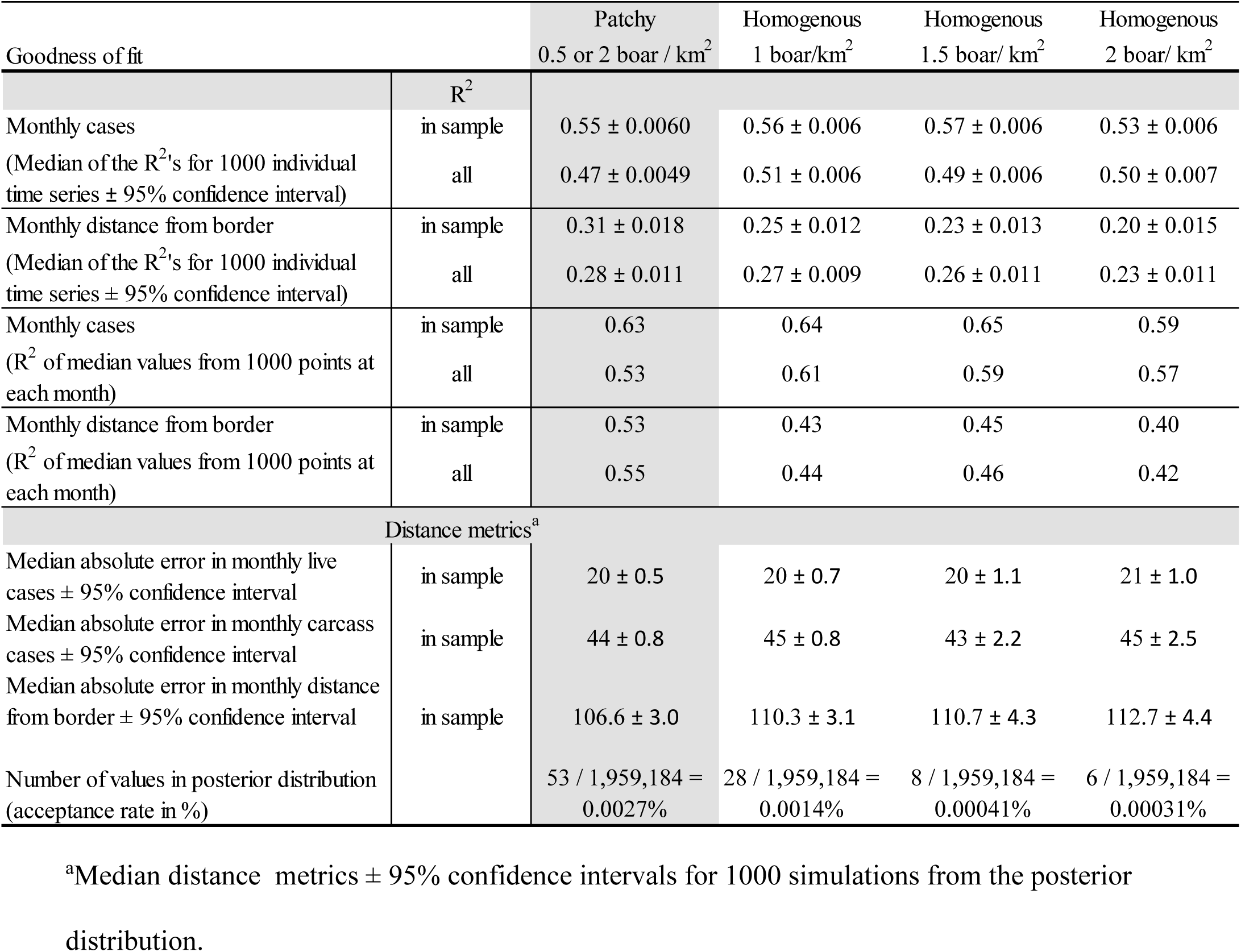
Model selection and fits for different densities and patchiness of the wild boar population. The model that best explained the data is highlighted in light grey. This model produced similar fits to the case data compared with the other models, but performed significantly better in terms of the R^2^ and ABC distance metrics for fitting to the distance from the border.

| Goodness of fit |  | Patchy<br>0.5 or 2 boar / km<sup>2</sup> | Homogenous<br>1 boar/km<sup>2</sup> | Homogenous<br>1.5 boar/ km<sup>2</sup> | Homogenous<br>2 boar/ km<sup>2</sup> |
| --- | --- | --- | --- | --- | --- |
|  | R<sup>2</sup> |  |  |  |  |
| Monthly cases<br>(Median of the R<sup>2</sup>'s for 1000 individual<br>time series ± 95% confidence interval) | in sample<br>all | 0.55 ± 0.0060<br>0.47 ± 0.0049 | 0.56 ± 0.006<br>0.51 ± 0.006 | 0.57 ± 0.006<br>0.49 ± 0.006 | 0.53 ± 0.006<br>0.50 ± 0.007 |
| Monthly distance from border<br>(Median of the R<sup>2</sup>'s for 1000 individual<br>time series ± 95% confidence interval) | in sample<br>all | 0.31 ± 0.018<br>0.28 ± 0.011 | 0.25 ± 0.012<br>0.27 ± 0.009 | 0.23 ± 0.013<br>0.26 ± 0.011 | 0.20 ± 0.015<br>0.23 ± 0.011 |
| Monthly cases<br>(R<sup>2</sup> of median values from 1000 points at<br>each month) | in sample<br>all | 0.63<br>0.53 | 0.64<br>0.61 | 0.65<br>0.59 | 0.59<br>0.57 |
| Monthly distance from border<br>(R<sup>2</sup> of median values from 1000 points at<br>each month) | in sample<br>all | 0.53<br>0.55 | 0.43<br>0.44 | 0.45<br>0.46 | 0.40<br>0.42 |
| Distance metrics<sup>a</sup> |  |  |  |  |  |
| Median absolute error in monthly live<br>cases ± 95% confidence interval | in sample | 20 ± 0.5 | 20 ± 0.7 | 20 ± 1.1 | 21 ± 1.0 |
| Median absolute error in monthly carcass<br>cases ± 95% confidence interval | in sample | 44 ± 0.8 | 45 ± 0.8 | 43 ± 2.2 | 45 ± 2.5 |
| Median absolute error in monthly distance<br>from border ± 95% confidence interval | in sample | 106.6 ± 3.0 | 110.3 ± 3.1 | 110.7 ± 4.3 | 112.7 ± 4.4 |
| Number of values in posterior distribution<br>(acceptance rate in %) |  | 53 / 1,959,184 =<br>0.0027% | 28 / 1,959,184 =<br>0.0014% | 8 / 1,959,184 =<br>0.00041% | 6 / 1,959,184 =<br>0.00031% |

<sup>a</sup>Median distance metrics ± 95% confidence intervals for 1000 simulations from the posterior distribution.

Rejection rates for the proposed parameter sets were high for all four models, such that posterior distributions ranged between 6-53 values (0.00031%-0.0027% model acceptance rate) (Table 2), and uncertainty in parameter estimates were large (See Fig. S6). Due to the high amount of stochasticity in model processes and uncertainty in parameter estimates, the model fit the data *on average* (i.e., R^2^ for the median trajectory of stochastic runs relative to observed data; Fig. 3c, d) better than the observed data relative to any one trajectory (i.e., median of R^2^’s for each stochastic run; Fig. 3a, b). R^2^ for the full data (including out-of-sample predictions) were lower than those for the in-sample predictions (Table 2, Fig. 3a, b), indicating that the model performed worse at out-of-sample prediction. The posterior distributions revealed parameter correlations (Fig. S7). β_d_ and β_c_ were negatively correlated with each other and even more negatively correlated with λ, whereas ρ_c_ and π were positively correlated (Fig S7). Other parameters were relatively uncorrelated.

### 3.2 Role of carcass-based transmission

The models predicted a substantial amount of carcass-based transmission (monthly average between 53 and 66% during 2014-2015 depending on the landscape; Fig. 4) and a much higher prevalence of ASF in sampled carcasses versus hunter harvested samples (Fig. S5a). The best model (patchy landscape) also predicted a slow decline of the wild boar population over time (Fig. S5b), which corresponded to proportionately more transmission events originating from carcass-based transmission over time (Fig. 4), especially in the patchy and low-density homogenous landscapes.

**Fig. 4.**
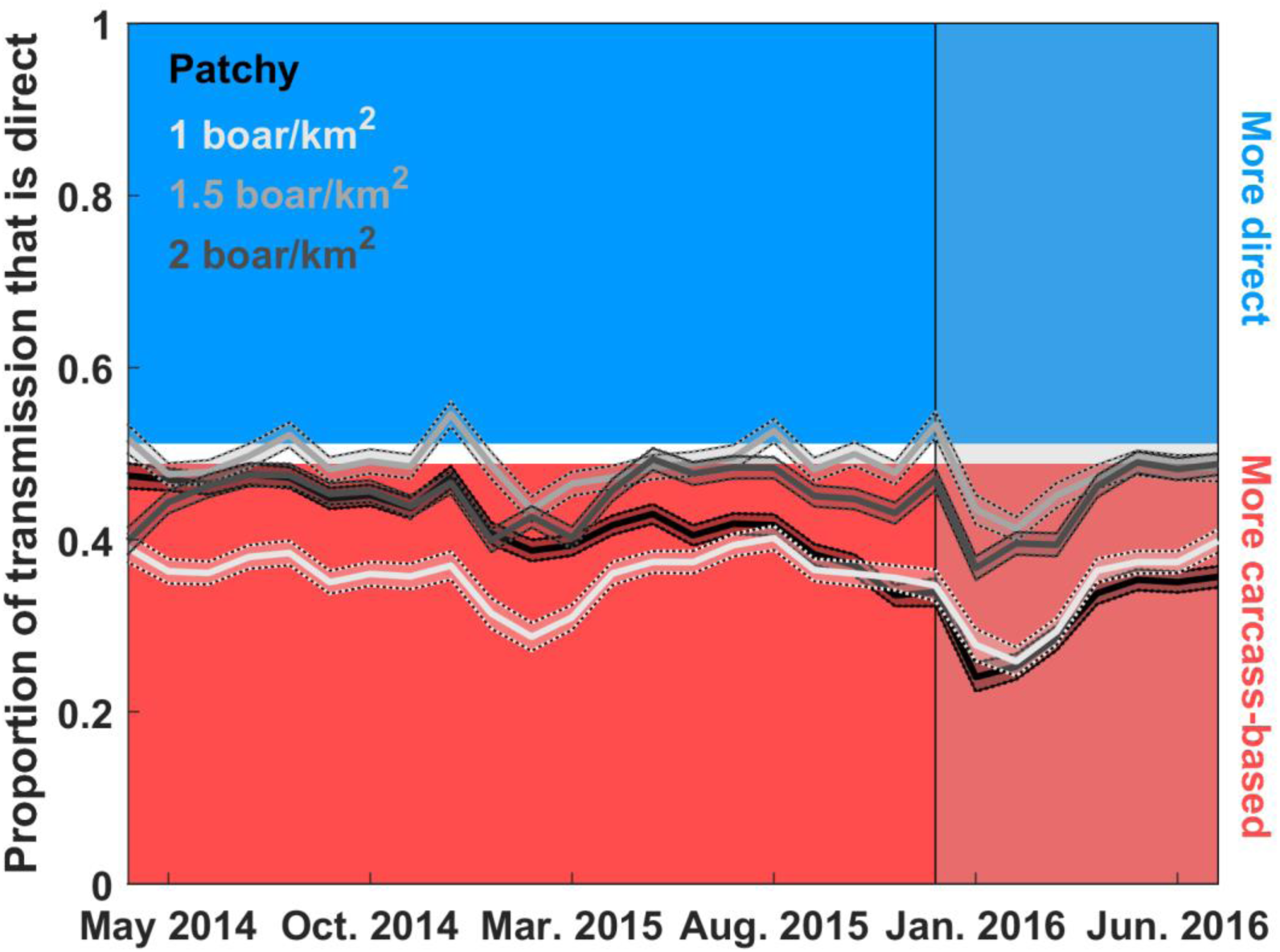
Proportion of transmission events that are from direct transmission. Shaded lines are 95% prediction intervals for 1000 simulations from posterior distributions of each model. Red indicates >50% of transmission events are carcass-based; blue indicates that >50% are direct. The different lines show results for different landscapes (heterogeneous versus the three homogenous landscapes of different densities). Note, parameter estimates were different depending on the landscape (Table 2). The transparent shaded panel indicates out-of-sample predictions.

### 3.3 Host density effects on ASF persistence

Sensitivity analyses showed that densities higher than 1 boar / km^2^ were important for autonomous persistence (Fig. 5 a,d,g). Without carcass-based transmission, persistence required re-introduction 10 or more times per year at lower host densities (Fig. 5b,e). However, with only carcass-based transmission, persistence occurred across some narrow range of carcass-based transmission probabilities even at low host densities (Fig. 5c,f) with few to no re-introductions. In contrast, high host density (4 boar / km^2^) allowed for autonomous persistence when carcass-based transmission was absent (Fig. 5g,h,i) over some narrow range of transmission probabilities for either transmission mechanism on its own.

**Fig. 5.**
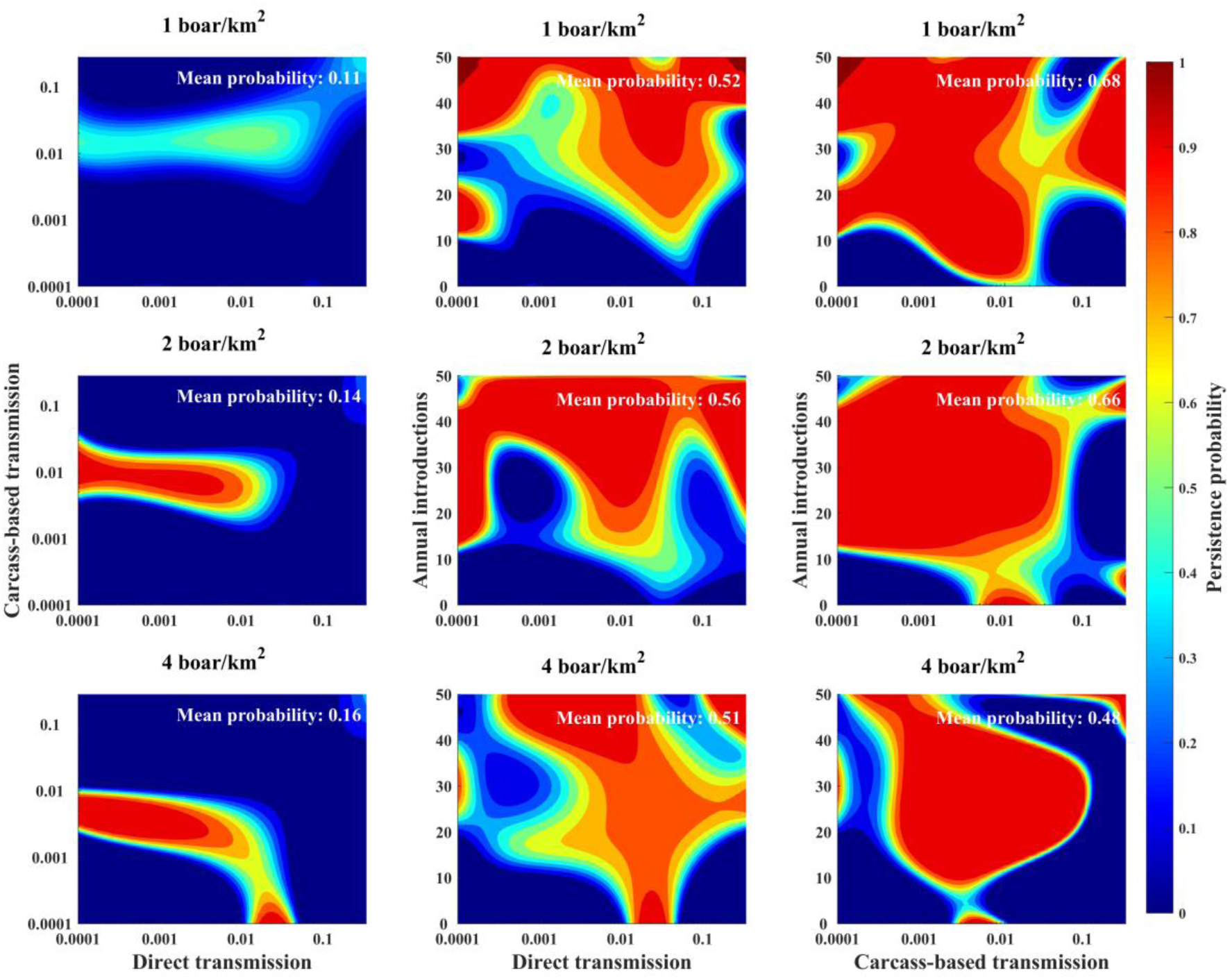
Effects of host density and persistence mechanisms. Colors show the probability that ASFV will persist with dark red representing high probability, yellow representing moderately high probability, light blue representing moderate probability, and dark blue representing low probability. Axes show the values of the three persistence processes we examined: 1) between-group direct transmission probability (β_d_), 2) between-group carcass-based transmission probability (β_c_), and 3) introduction frequency (ϕ) as indicated. For each two-way plot, the third parameter (β_d_, β_c_, or ϕ) was fixed at 0 in order to disentangle each two-way interaction. Within-group transmission probabilities were fixed at 10 times their respective between-group transmission probabilities. Other parameters estimated by the model were fixed at biologically realistic values: ρ_h_ = 0.015, ρ_c_ = 0.025, π = 1, θ = 2, λ = 1.5; other parameters were as in Table 1. Each plot shows results for a different host density. The mean values in black show means for the entire plot, giving an overall effect of the landscape on spatial spread.

## 4 DISCUSSION

The persistence of ASFV in wild boar in Eastern Europe remains a significant threat to domestic pig populations globally, and hence international trade and food security. By fitting a mechanistic disease-dynamic model to spatio-temporal disease surveillance data using prior knowledge of wild boar population dynamics, we inferred that 53-66% of ASFV transmission events occurred through the contact of susceptible hosts with dead carcasses. Because wild boar tend not to contact carcasses immediately, but will continue to contact carcasses even during the later stages of decay (Probst et al. 2017), increased surveillance and elimination of carcasses could dramatically decrease transmission (Morelle et al. 2019). Thus, developing cost-effective methods for carcass detection and retrieval may be critical to reduce transmission rates in wild boar populations (Guinat et al. 2017). Additionally, as we found that the relative importance of transmission mechanisms depended critically on host density, our results emphasize the importance of considering wild boar population dynamics in control (EFSA, 2017).

Although our model captured the data fairly well, when the model fit incidence as well as possible, it underestimated the rate of spatial spread. When it fit the rate of spatial spread well, it overestimated incidence. Thus, the structure of our model lacked an important unknown process in the spread of ASFV in wild boar populations. Because our modeling framework accounts for well-documented spatial contact (Podgórski et al. 2018) and dispersal distances (Keuling et al. 2010, Podgórski et al. 2014b), our difficulty with concurrently fitting incidence and distance trajectories could be due to long-distance movements occurring often enough to seed infection outside the daily home range or dispersal movements. These results come as no surprise, as the human-mediated spread of ASFV continues to play a large epidemiological role in the area (EFSA, 2017). One possible source is hunters that may contaminate hunting equipment when processing infectious carcasses and then introduce the infectious fomites at another site (Wieland et al. 2011). Implementing enhanced hunting biosecurity policies could help reduce the spatial spread of ASFV by hunters. A second possibility are other species which have contact with infectious carcasses, e.g. scavenging carnivores, birds, and flies, and subsequently disperse contaminated tissue. Although the role of mechanical vectors in ASFV epidemiology remains unknown, they are known to enhance spread of many diseases (Siembieda 2011), suggesting that this mechanism could be worth further investigation.

Our model tended to predict a relatively high incidence immediately after introduction of the index case, as would be expected by a new disease introduced into a completely susceptible population. However, hunting was limited initially in the 1^st^ 3-4 months which accounts for the low number of observed cases during this time. Once the hunting ban was lifted, the number of observed cases increased, although the underlying dynamics were more consistent (data not shown). We accounted for temporal trends in hunting using data on overall sample sizes, but we did not have precise locations for negative surveillance data. Locations of all samples are important for data fitting because, as we saw with the temporal sampling data, the surveillance system limits our observation of the underlying process. If in reality surveillance sampling locations shifted in spatial clusters that were nearer versus farther from the border, rather than all samples being spread out randomly, it is possible that cases that were further from the border may have been detected – i.e., representing the surveillance process as spatially random could have diluted detection of cases that occurred further away. Indeed, the true rates of spatial spread were faster than the predicted observed rates in our model (e.g., 16.8 vs 20.2 km from the border respectively for year 1, and 26.8 vs 31.5 km respectively for year 2), suggesting that accounting for spatial locations of all surveillance samples could help to improve inference of the spatial spreading process, and potentially our understanding of the role of transmission mechanisms. Our analyses highlight the importance of fully recording metadata for negative samples and appropriately accounting for the full sampling design in the observation process.

Several studies have observed reductions in prevalence of target pathogens from within populations that diseased hosts are removed from (e.g., Donnelly et al. 2006, Mateus-Pinilla et al. 2013, Manjerovic et al. 2014, Boadella et al. 2012). However, as others have emphasized, there can be unexpected consequences of culling programs – for example, adaptation of the virus to low density conditions (Bolzoni & De Leo 2013), or increases in long-range host movements that lead to increased spread of disease (Bielby et al. 2014, Comte et al. 2017). Our sensitivity analyses also demonstrated that decreasing host density could have unexpected consequences. We found that while high host densities allowed autonomous persistence by direct transmission, carcass-based transmission could allow persistence as host density decreases by effectively extending the opportunity for hosts to contact contaminated carcasses. Thus, a thorough understanding of the host-pathogen transmission ecology in response to management are important before planning abundance reduction programs (Harrison et al. 2010). To control ASFV, density reduction programs will likely be most successful if they include intensive surveillance and the removal of dead carcasses, especially as populations reach low densities.

A recent study found that for wild boar, a social species that aggregates in family groups (Podgórski et al. 2014a, Podgórski et al. 2014b), spatially-targeted culling that focuses on removing all members of family groups is more effective than random removal of individuals (Pepin & VerCauteren 2016). Although this simulation model did not account for movement responses due to culling, it did include effects of social structuring in disease transmission. Because intensive hunting can induce escape movements and intergroup mixing (e.g., Scillitani et al. 2010), culling programs should consider removal of all individuals in a group to decrease the chance of increased long-distance movement due to social structure disruptions.

We assumed that ASF was 100% lethal in wild boar, which is an oversimplification of the system. Indeed, surveillance data in the region suggest there are survivors with < 1% of individuals testing positive for antibodies against ASFV in the outbreak region (unpublished data of the National Veterinary Research Institute, Puławy, Poland). Because we assumed absolute lethality, our models predicted that ongoing transmission of ASFV led to decreased host densities and higher levels of carcass-based transmission over time. However, if a strain of ASFV with reduced lethality were to emerge in this area we might expect different epidemiological dynamics. Specifically, if more hosts remained alive, they would be available to reproduce, maintaining higher host densities and a more consistent influx of naïve hosts which could lead to persistence or recurrent epidemics by direct transmission alone (Stone et al. 2007). Thus, surveillance for changes in lethality are important for optimizing control strategies in local areas over time.

Our implementation of wild boar spatial processes is a simplification of reality. We assumed that individuals could potentially contact other individuals in all directions each day, with a contact probability that decayed with distance. However, in reality, wild boar movements are biased towards habitat features and related individuals (Kay et al. 2017, Podgórski et al. 2014a). Although these movements would lead to temporal variation in the distance-transmission probability relationship as we assumed, it may be that the particular locations of the movements (e.g., biased towards particular resources or conspecifics) are important to capture, rather than just the overall distance-variation structure. Indeed, recent work has shown that accounting for elk movement mechanistically can provide predictions of spatio-temporal prevalence of brucellosis disease in response to changing seasonality and climate (Merkle et al. 2018). Mechanistic movement models based on landscape heterogeneity have also been used to predict areas of highest contact rates (Tardy et al. 2018), another metric of disease transmission risk. Adaptive prediction of where and when disease transmission risk may be highest is important for enabling managers to prioritize mitigation strategies in space and time in a cost-effective manner. Additionally, although theory predicts that implementing reactive control based on knowledge of social structure can help determine the effectiveness of control (e.g., Azman et al. 2015), we rarely know individual-level relationship status *a priori*. Understanding the link between mechanistic movement, landscape heterogeneity, and disease transmission could help provide more practical (landscape-based) guidance for prioritizing surveillance and interventions.

## ACKNOWLEDGEMENTS

KMP was supported by the United States Department of Agriculture, Animal and Plant Health Inspection Service’s National Feral Swine Damage Management Program. AG was supported by the United States Department of Agriculture, Animal and Plant Health Inspection Service’s APHIS Science Fellowship. TP was supported by the National Science Centre, Poland (grant number 2014/15/B/NZ9/01933). We thank M. Łyjak, A. Kowalczyk, K. Śmietanka, and G. Woźniakowski from Department of Swine Diseases, National Veterinary Research Institute in Pulawy, Poland, for surveillance data. N. Selva provided valuable information on carcass persistence time.

## DATA ACCESSIBILITY

Data will be archived at the National Wildlife Research Center data archive and Dryad.

## AUTHORS’ CONTRIBUTIONS

KMP and TP conceived the idea. KMP, TP, and ZA contributed to model development. KMP performed the analyses. KMP led the writing. All authors contributed to critical analysis of results, manuscript editing, and gave final approval for publication.

## Supplementary Figures

**Fig. S1.**
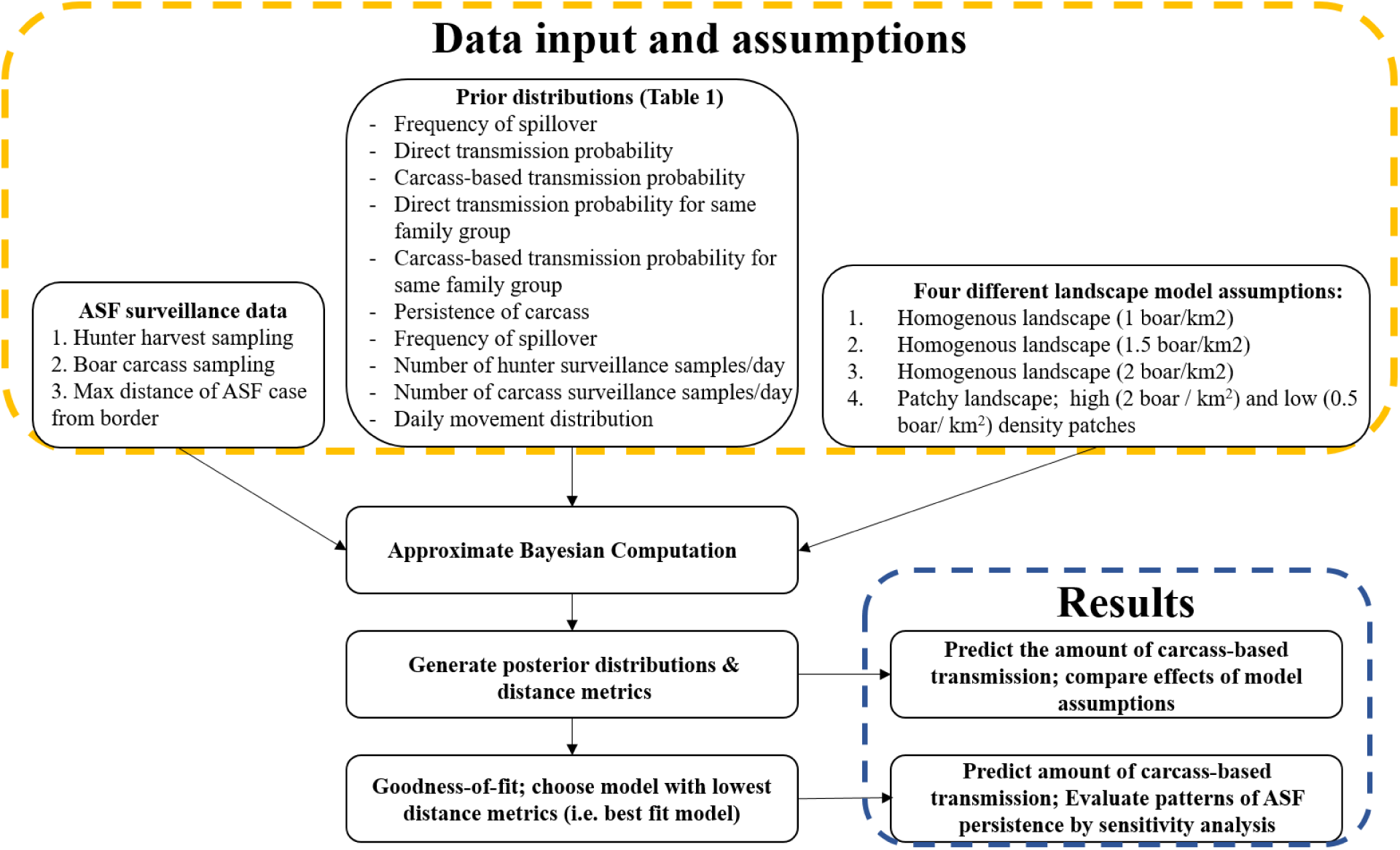
**Schematic of modeling approach.**

**Fig. S2.**
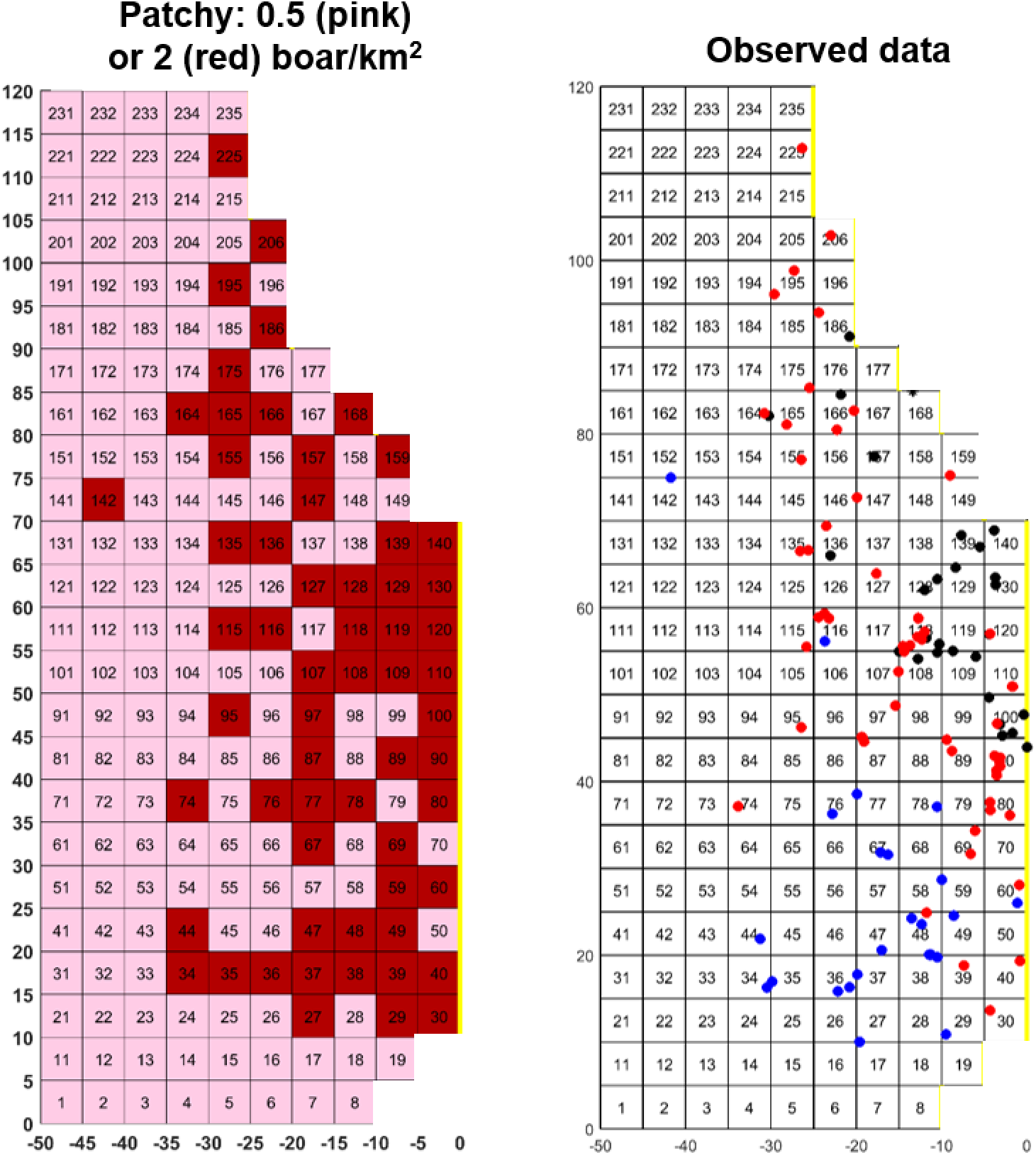
**Host density structure of landscapes used for data fitting by ABC.** X and Y-axes are in km. The Eastern border is indicated in yellow – boar were not allowed to the right of this line. Each grid cell is 5 x 5 km. Grid cell colors correspond to maximum boar density allowed in the grid cell which was 0.5 (light pink) or 2 (dark red) boar / km^2^. This landscape was the patchy landscape. We also compared fits on this landscape to ones with homogenous densities of 1 (the average of the patchy landscape), 1.5, and 2 boar/km_2_ to evaluate the role of landscape structure in explaining the patterns of spread. The grid on the right shows observed case locations by year: 2014 (black), 2015 (red), Jan.-Jul. 2016 (blue). The data from 2016 was withheld for parameter estimation, but was used to assess out-of-sample model performance.

**Fig. S3.**
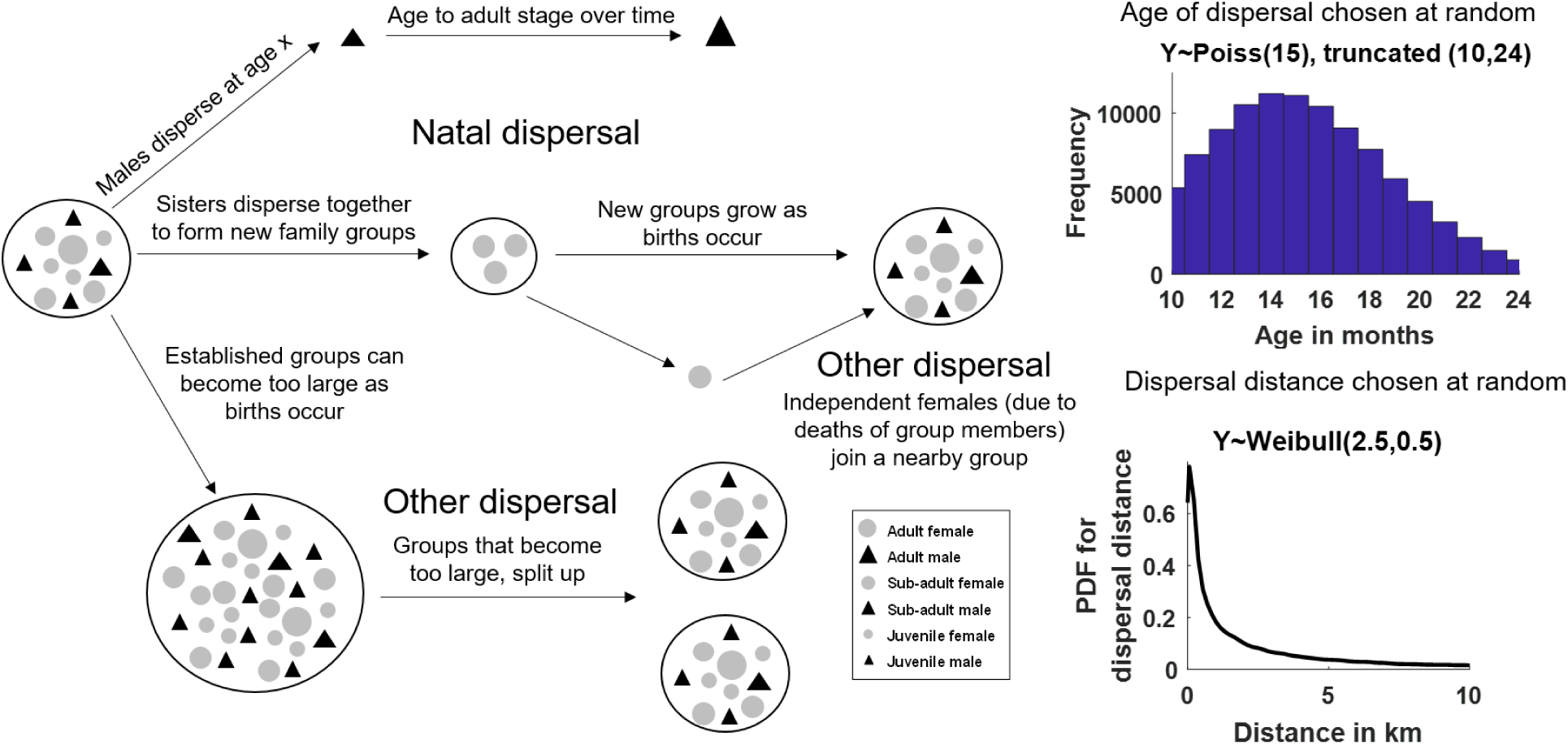
**Dispersal and social dynamics.** Schematic of dispersal mechanisms and social dynamics. These dispersal events result in permanent relocation of the home range centroid. Natal dispersal age and dispersal distance for each permanent relocation event are chosen at random from distributions supported by empirical data displayed on the right half of the figure.

**Fig. S4.**
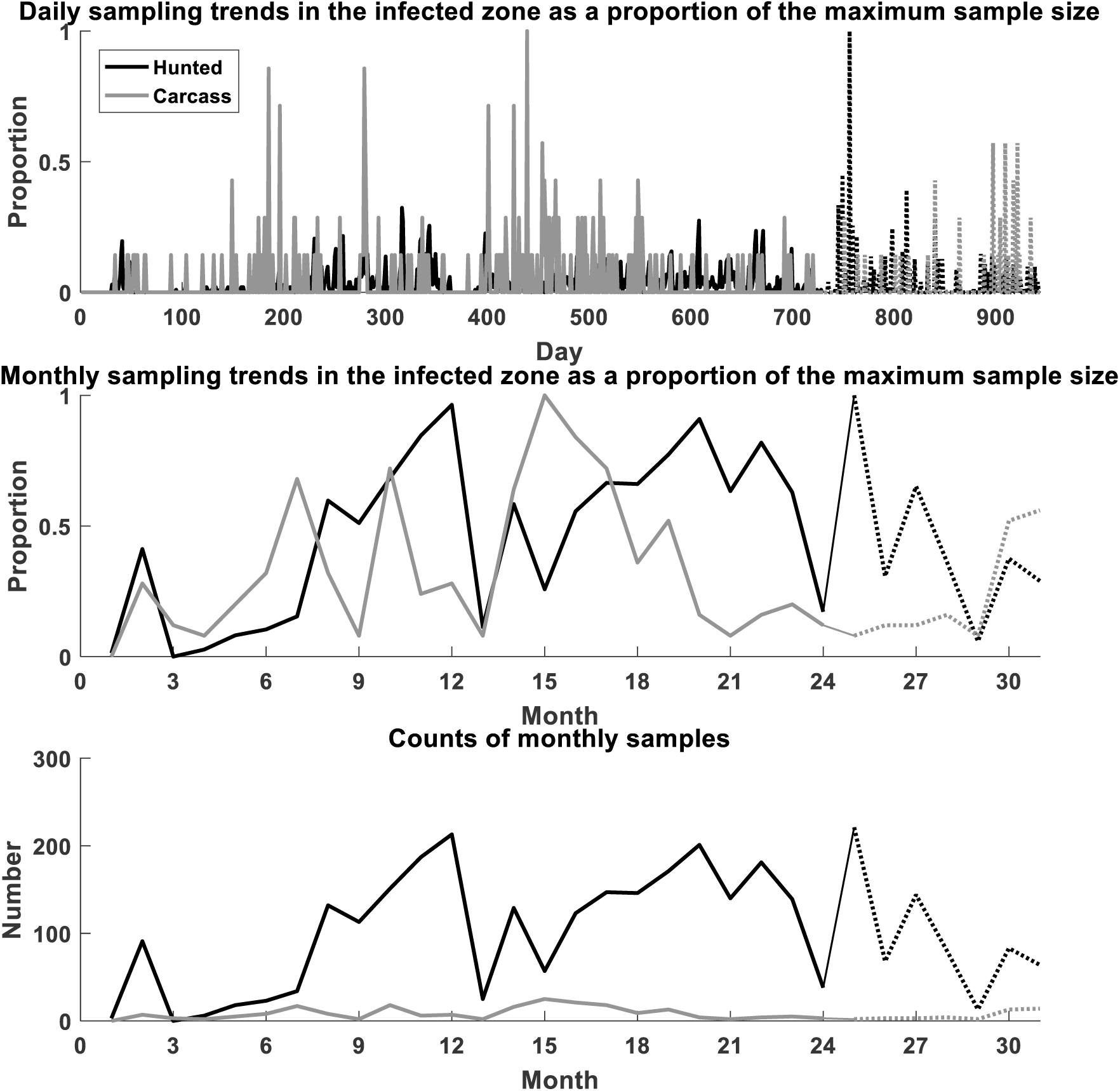
**Sampling trends in the infected zone.** The vectors of daily proportion values were multiplied by scaling parameters (ρ_h_, ρ_c_) to determine the proportion of the population sampled each day. Thus, for example, the number of boar sampled on day *t* by hunting was: number of boar on the landscape (only including those > 6 months) x proportion in the sampling trend vector on day *t* x ρ_h_. This method accounts for seasonal changes in boar abundance and sampling. The dotted lines indicate 2016 data, which were not used for parameter estimation but were used for prediction out-of-sample.

**Fig. S5.**
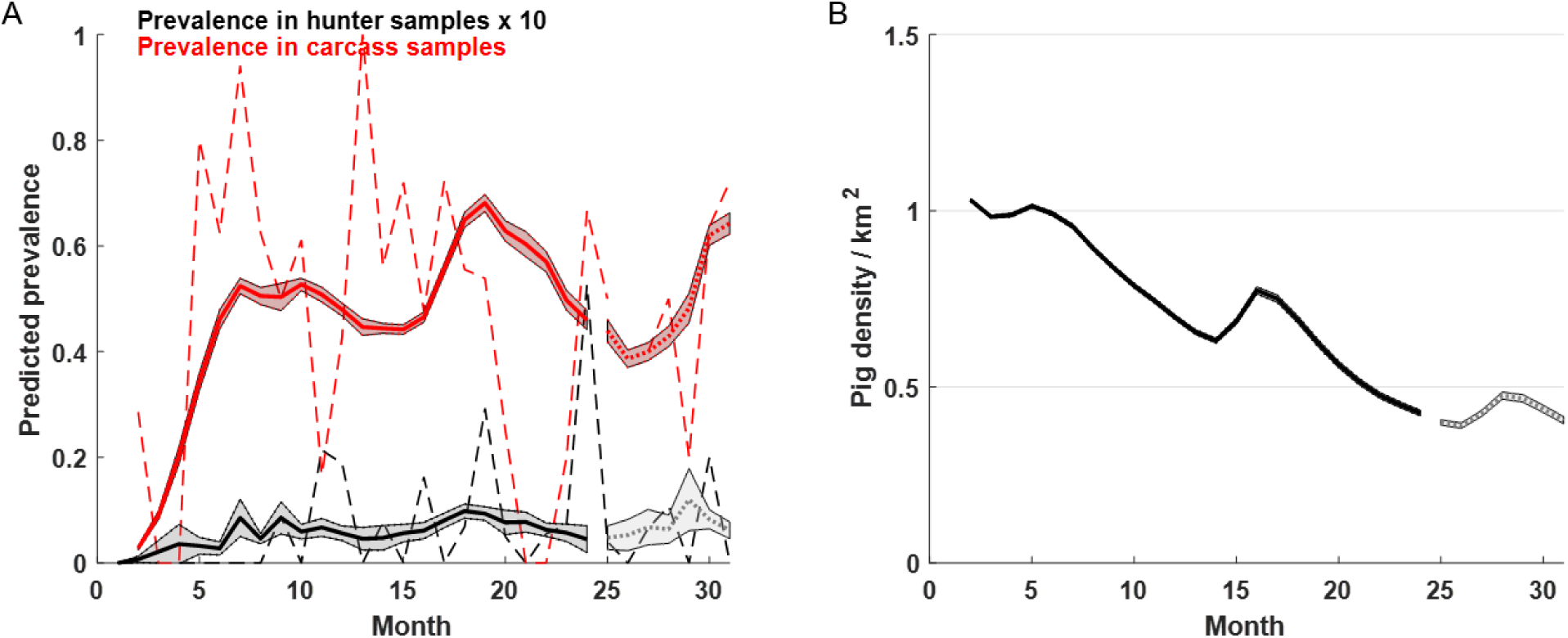
Predictions from 1000 simulations from the posterior distribution (patchy landscape). Solid lines indicate the time period used for model fitting whereas dotted lines show the out-of-sample predictions for 2016. A. Monthly prevalence in hunted (black) and carcass (red) surveillance samples. Dashed lines are monthly prevalence in the real surveillance data. B. Predicted abundance of wild boar in the simulations.

**Fig. S6.**
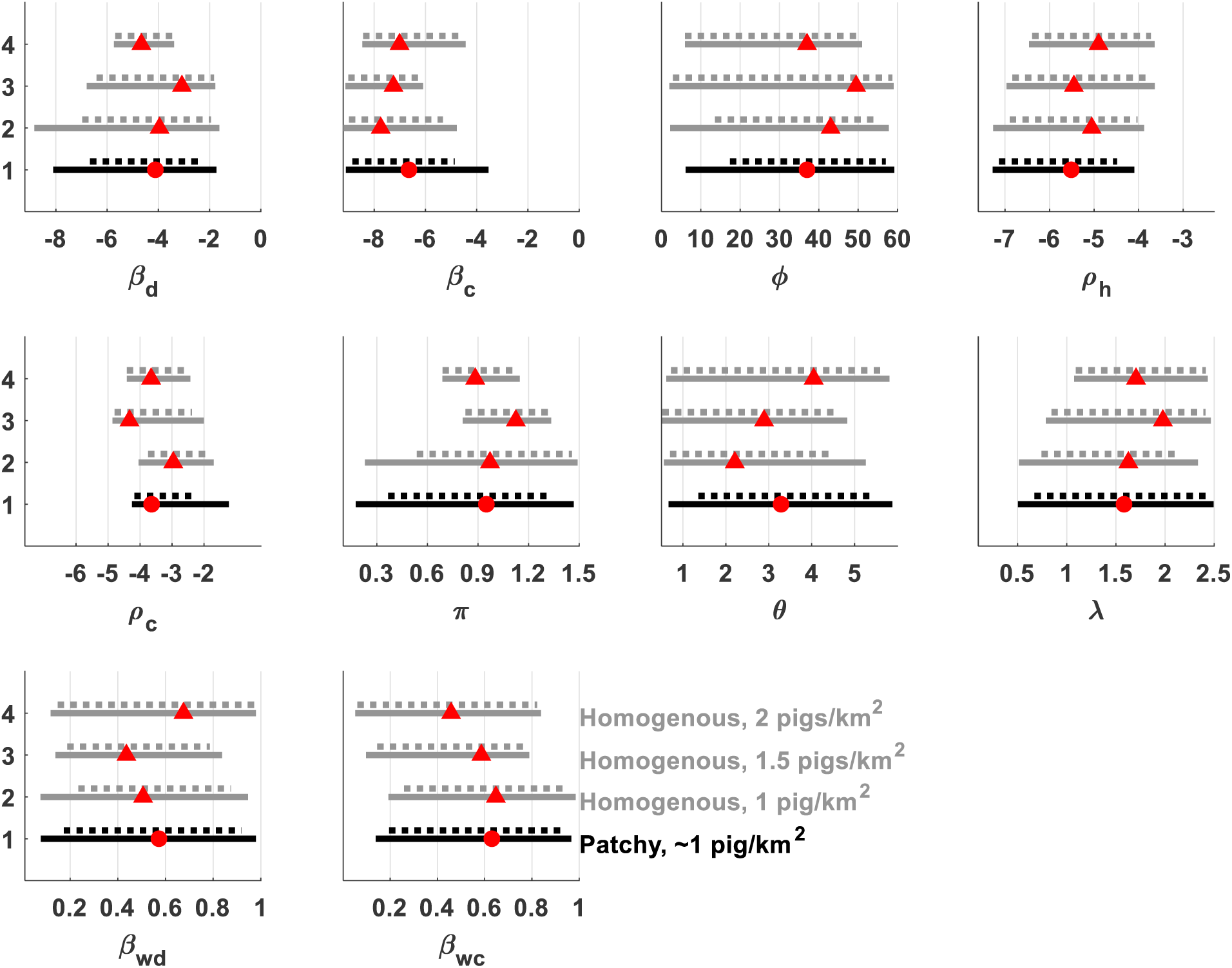
Credible intervals (CI) of estimated parameters for each model (solid lines are 95% CIs, dotted lines are 80% CIs). Prior distributions are all uniform and span the width of the X-axes. Red shapes indicate the median of the posterior distribution.

**Fig. S7.**
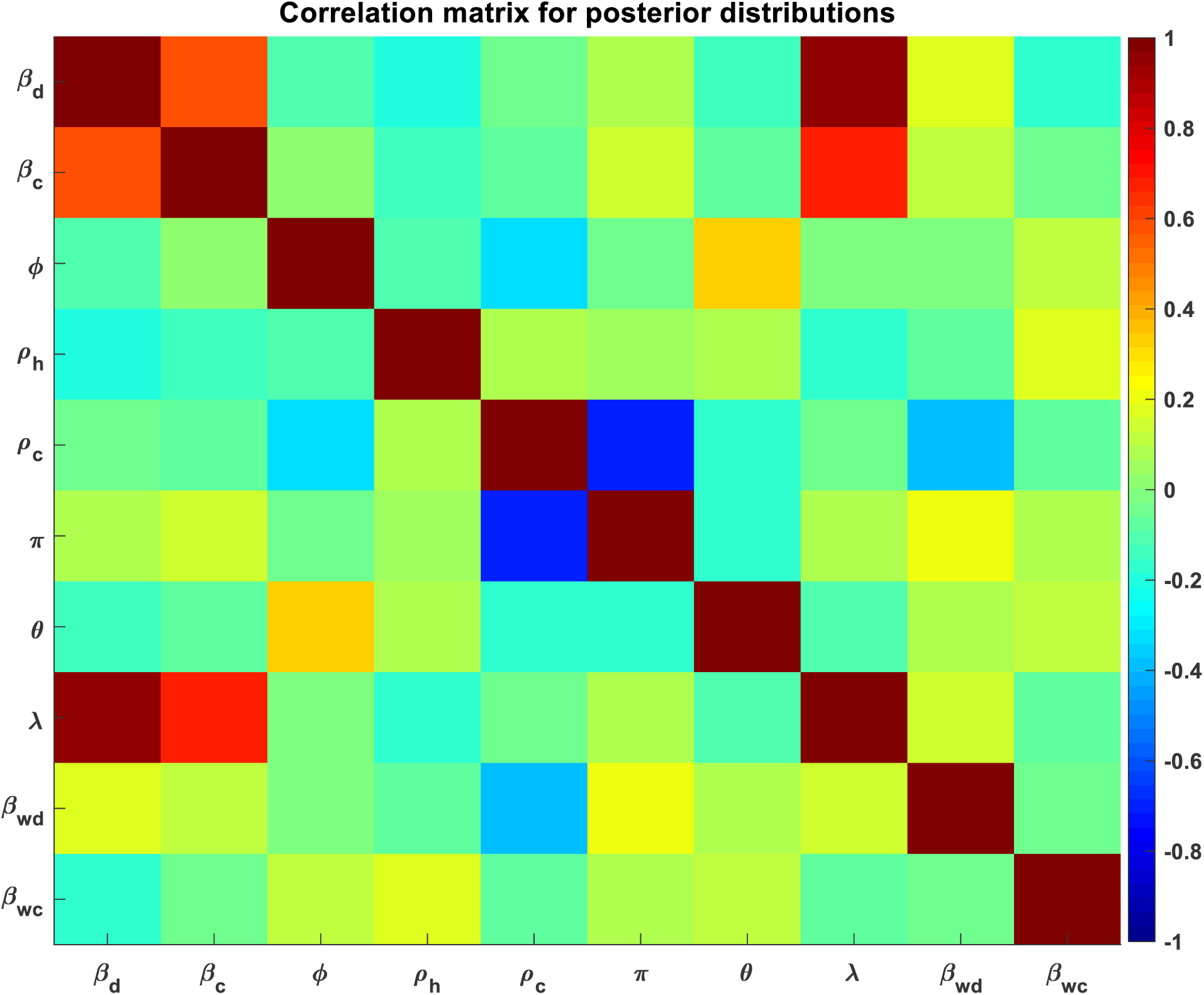
Results are from the patchy landscape. Colors are correlation coefficients between parameters. β_d_ and β_c_ were positively correlated with each other and even more positively correlated with λ, whereas ρ_c_ and π were negatively correlated, and other parameters were relatively uncorrelated.

